# Improved prediction of bacterial CRISPRi guide efficiency from depletion screens through mixed-effect modeling and data integration

**DOI:** 10.1101/2022.05.27.493707

**Authors:** Yanying Yu, Sandra Gawlitt, Lisa Barros de Andrade e Sousa, Erinc Merdivan, Marie Piraud, Chase L. Beisel, Lars Barquist

## Abstract

CRISPR interference (CRISPRi), the targeting of a catalytically dead Cas protein to block transcription, is the leading technique to silence gene expression in bacteria. However, design rules for CRISPRi remain poorly defined, limiting predictable design for gene interrogation, pathway manipulation, and high-throughput screens. Here we develop a best-in-class prediction algorithm for guide silencing efficiency by systematically investigating factors influencing guide depletion in multiple genome-wide essentiality screens, with the surprising discovery that gene-specific features such as transcriptional activity substantially impact prediction of guide activity. Accounting for these features as part of algorithm development allowed us to develop a mixed-effect random forest regression model that provides better estimates of guide efficiency than existing methods, as demonstrated in an independent saturating screen. We further applied methods from explainable AI to extract interpretable design rules from the model, such as sequence preferences in the vicinity of the PAM distinct from those previously described for genome engineering applications. Our approach provides a blueprint for the development of predictive models for CRISPR technologies where only indirect measurements of guide activity are available.

## Introduction

CRISPR interference (CRISPRi), in which a catalytically dead Cas protein incapable of DNA cleavage (dCas) is targeted to interfere with transcription of a gene of choice [1,2], is one of most widely used CRISPR technologies in bacteria. In contrast to eukaryotes, many bacteria lack the necessary repair pathways to survive genome editing by the double-stranded break induced by CRISPR-Cas9. Thus, CRISPR-Cas’s main applications in bacteria have come from using dCas as a platform technology that can deliver effectors to a specific locus in a programmable fashion. CRISPRi is the simplest example, where the dCas protein itself serves as an effector to silence gene expression by physically blocking the binding or procession of the RNA polymerase.

The development of CRISPRi has opened up a range of biological applications, from silencing individual genes for genetic studies to performing genome-wide fitness screens or engineering genetic circuits [3,4]. As an alternative screening technology to transposon mutagenesis [5], CRISPRi can directly target particular genes of interest, avoiding the need for large mutant libraries to achieve gene saturation. Another area of application is engineering synthetic regulatory circuits [6] or metabolic networks [7,8], where collections of gRNAs are used to coordinately downregulate and upregulate associated genes and pathways. However, all of these applications critically depend on the efficiency of silencing provided by selected guides. Genetic screens already routinely employ tens of thousands of guides simultaneously, and it is impractical to individually test each guide’s efficiency. This problem will only be accentuated as the scale of applications increases through the use of CRISPR array technologies that allows multiplexed expression of suites of guides simultaneously [9,10] to dissect and engineer increasingly complex phenotypes. Reliable prediction of guide efficiency will therefore become increasingly important as applications of CRISPRi become more ambitious.

Given the impact of CRISPR-based genome engineering in eukaryotes, significant effort has been expended in developing methods for predicting editing efficiency. The first attempts used classical machine learning methods on relatively small datasets comprising efficiency measurements for thousands of gRNAs. The applied methods include logistic regression [11], support vector machines [12,13], linear regression [14], and gradient-boosted decision trees [15]. As the amount of Cas9 editing data has increased, deep learning approaches have become increasingly popular. These approaches include convolutional neural networks [16], which apply a collection of adaptive filters to automatically extract local sequence features, and long short-term memory networks (LSTM) [17], which retain a memory that potentially allows for the detection of long-range interactions between sequence features. Newer methods have put substantial effort into engineering deep learning architectures to further boost performance [18]. It is important to note that many of these deep learning methods have been trained on tens of thousands of measurements of guide efficiency, and fusing datasets has played an important role in further increasing performance [19].

So far, relatively little attention has been paid to predicting guide efficiency for bacterial CRISPRi. The only study to date developed a LASSO regression model for predicting CRISPRi guide efficiency [20] with a limited sequence feature set using data from a single genome-wide CRISPRi screen in *Escherichia coli* [21], providing important proof-of-concept but also leading us to ask if larger datasets and more complex machine learning approach could further improve prediction. Here, starting with an investigation of the features driving depletion of guides targeting essential genes in CRISPRi screens using automated machine learning, we find that features associated with the targeted gene, such as expression, explain most of the variation in the data. Starting from this foundation, we develop a mixed-effect random forest that separates features affecting guide efficiency from effects due to the targeted gene while learning from multiple independent CRISPRi screens, allowing us to arrive at a predictive model of guide efficiency that we show improves on the state-of-the-art using a saturating depletion screen of purine biosynthesis genes during growth in minimal medium. Our mixed-effect machine learning approach provides a general strategy for learning CRISPRi guide efficiency when only indirect measurements are available.

## Results

### Automated machine learning and feature engineering identifies gene-specific effects in CRISPRi depletion screens

We set out to devise design rules for CRISPRi in bacteria by combining machine learning with large experimental datasets. The largest available datasets come from genome-wide depletion screens which provide only indirect measurements of guide efficiency (**Figure 1A**). We began our investigation into the features affecting guide depletion by applying automated machine learning (autoML) in the form of the Auto-Sklearn package [22] that wraps classification and regression models implemented in the Python Scikit-learn package [23] in a Bayesian optimization framework.

**Figure 1:**
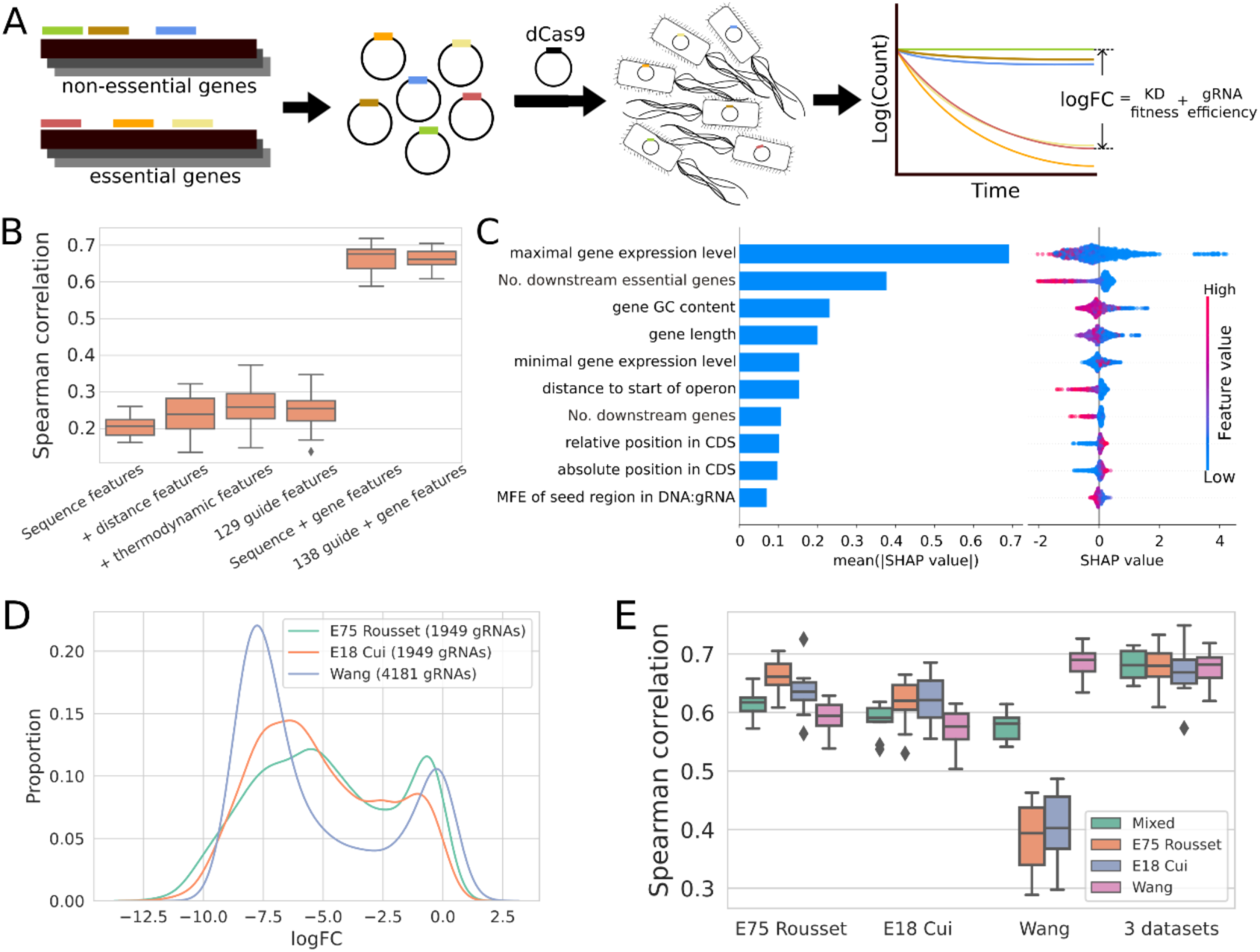
Automated machine learning and data fusion predicts depletion in CRISPRi essentiality screens. (**A**) An overview of CRISPRi essentiality screens. gRNAs are designed targeting every gene in the genome and cloned into an appropriate plasmid for expression. This plasmid collection is then transformed into the target bacteria, and depletion is measured as the change in guide frequency over growth determined by sequencing relative to a set of non-targeting gRNAs. The measured depletion (logFC) is then a mixture of the fitness effect of gene knockdown with the efficiency of silencing itself. **(B)** Comparison of Spearman correlation between actual and predicted guide depletion in 10-fold cross-validation (CV) of the best model trained with Auto-Sklearn with different feature combinations, using data from [21]. **(C)** The ten most predictive features determined using TreeExplainer on the optimal random forest model trained with Auto-Sklearn and 574 guide and gene features. Mean absolute SHAP value (left) provides a global measure of feature importances, while the beeswarm plot (right) shows the effect of each feature on each individual gRNA prediction. CDS: coding sequence. (**D**) Distribution of logFCs of gRNAs targeting essential genes from three CRISPRi genome-wide essentiality screens in *E. coli*. (**E**) Comparison of Spearman correlation from 10-fold CV of the best Auto-Sklearn trained model on one dataset or the three integrated datasets.

We first asked how well we could predict gRNA depletion log_2_ fold-changes (logFCs) for essential genes as defined by the Keio collection [24], and what features would be required for accurate prediction. As essential genes should have an infinite fitness cost upon complete silencing, we initially assumed differences in depletion would mainly depend on features describing the gRNAs. We leveraged a published *E. coli* CRISPRi essentiality screen using dCas9 performed in rich medium [21], which included 1,951 guides targeting 293 essential genes. To predict depletion in this dataset, we engineered a series of feature sets of increasing complexity (**Figure S1**; **Table S1**) starting with the one-hot-encoded gRNA and PAM sequence including four bases upstream of the gRNA target sequence and three bases following the NGG PAM. This resulted in a poorly performing model with a median Spearman’s ρ of ∼0.21 in 10-fold cross-validation (**Figure 1B; Table S2, S3**). We therefore iteratively added a set of additional features while monitoring changes in model performance. As targeting efficiency in bacteria has been suggested to depend on distance to the transcriptional start site [1,25], the set included absolute and relative distance to the start codon. We also included a suite of thermodynamic features describing gRNA:target interactions predicted using the ViennaRNA package [26]: minimum free energy of the folded gRNA, hybridization of two gRNAs, and hybridization of the targeted DNA and gRNA [27]. These additional feature sets resulted in only moderate improvement in Spearman correlation (ρ ∼ 0.25) for our predictions.

Given that features describing the guide sequences themselves were inadequate to predict guide depletion, we developed a series of genomic and expression features describing each targeted gene to investigate if they could improve prediction of guide depletion (**Figure 1B**; see also Feature engineering, **Methods**; **Table S1**). We collected publicly available RNA expression data over growth in minimal medium [28] and computed minimum and maximum expression values. We collected transcription unit (TU) information from RegulonDB [29] and calculated the distance from the target site to the start of the TU, the number of downstream genes in each TU, and the presence of other essential genes in the TU, and gene GC content. Incorporating these additional gene features led to a major improvement in prediction accuracy, with cross-validation performance jumping to a Spearman’s ρ of ∼0.66. This shows that autoML can rapidly produce predictive models of CRISPRi guide depletion, given a sufficiently rich feature set describing both guide and gene features.

To understand the contribution of these features to the prediction of gRNA depletion, we used SHapley Additive exPlanation values (SHAP values) computed with TreeExplainer [30] on the best performing random forest model produced by Auto-Sklearn (**Figure 1C**; **Table S4**). Of all considered features, maximal RNA expression in minimal medium had the single largest average effect on depletion, making an average of an ∼1.6 fold-difference to the predictions. Unexpectedly, high target gene expression tended to be associated with higher depletion. The number of downstream essential genes also had a strong effect on depletion predictions with an average ∼1.3-fold difference, indicating the presence of polar effects in CRISPRi screens. The rest of the top seven features all described the targeted gene, including GC content, gene length, and the distance to the operon start, but each had smaller effects than expression. The most predictive effects that could be manipulated during guide design were associated with guide distance to the transcriptional start site, but on average these had fairly small effects (∼1.07 fold) compared to features associated with the target gene. These findings show that predictions made by our depletion models are dominated by the effects of gene features that can not be modified in guide design.

### Data fusion improves prediction of gRNA depletion

Given that several gene essentiality screens have now been performed in *E. coli*, we next asked whether expanding our training dataset could improve the accuracy of our predictions, as has previously been shown for eukaryotic CRISPR editing applications [19]. To this end, we collected data from two additional CRISPRi screens of *E. coli* in rich medium. First, we included data from an additional screen using the same gRNA library but with dCas9 expressed from a stronger promoter, which we refer to here as E18 Cui [31]. Second, we included data from a completely independent screen using a higher density library containing twice as many guides targeting essential genes (4,197, with 528 identical to gRNAs contained in Cui/Rousset), which we refer to as Wang [25]. We refer to the original data set as E75 Rousset. It is also worth noting that while the E18 Cui and E75 Rousset libraries were grown repeatedly to stationary phase, the Wang library was collected in log phase. The level of depletion in each dataset exhibited qualitative differences, with Wang showing a clearer bimodal separation between depleted and non-depleted guides (**Figure 1D**) and a reasonable correlation of depletion between datasets (ρ∼0.9 between E18 Cui and E75 Rousset; ρ∼0.75/0.79 between Wang and Cui/Rousset; **Figure S2**).

To investigate the impact of fusing these datasets on model performance, we trained a series of models using Auto-Sklearn with each dataset individually or in combination and then tested them on sets of guides held out from each dataset as well as a mixed test set (**Figure 1E**; **Table S5**). We additionally included a one-hot encoded dataset indicator as a predictor of potential batch effects. Unsurprisingly, models trained on single datasets tended to perform best on their cognate test set. Similarly, models trained on E18 Cui and E75 Rousset appeared to generalize better to each other than to the Wang dataset and vice versa. Combined training datasets produced models that generalized better across datasets (ρ ∼ 0.58—0.62 tested on mixed data, vs ρ ∼ 0.68) without degrading performance relative to models trained on individual datasets. In some cases, particularly for the Cui dataset, fused training sets actually improved performance on a test set drawn from a single dataset (ρ ∼ 0.62 trained on Cui; ρ ∼ 0.67 trained on fused data). In each case, the best performing model chosen by Auto-Sklearn was either a random forest regression or a gradient-boosted decision tree model. In support of this observation, we trained a suite of different regressors individually and found that while all models benefited from data fusion, the gains were particularly large for random forest and gradient boosted tree models (**Figure S3**). These findings suggest that both increased generalizability and accuracy can be achieved by integrating multiple data sources for training tree-based models for CRISPRi depletion.

### Segregating guide and gene effects produces a highly predictive model for CRISPRi guide efficiency

Our exploration of the features most predictive of gRNA depletion in competitive screens highlighted that features describing the targeted gene often made much larger contributions to the prediction than features describing the guide sequence. This is problematic for predicting guide efficiency from depletion screens, as this large gene-to-gene variation in depletion must be removed to properly extract the contribution of guide efficiency.

To isolate guide efficiency from measurements of depletion, we developed a method to fit and remove gene effects using Mixed-Effect Random Forest (MERF) regression [32] (**Figure 2A**). The MERF model handles data with an underlying cluster structure by defining two separate models: a linear model that captures random effects associated with the cluster, and a random forest (or other complex model) that captures fixed effects associated with each individual measurement after removing cluster effects. These models are then jointly optimized in an iterative process using the expectation-maximization algorithm. In our case, the cluster structure corresponds to collections of guides targeting the same gene. Random effects capture the effects of features associated with each gene (e.g. expression level) as well as dataset. Fixed effects are fit to the residual guide efficiency after removing the effects of the targeted gene, and correspond to features that could be manipulated in gRNA design (e.g. PAM and guide sequence, thermodynamic properties). The final fixed effect model can then be used to predict the relative efficiency of guides targeting a gene of interest independently of gene features.

**Figure 2:**
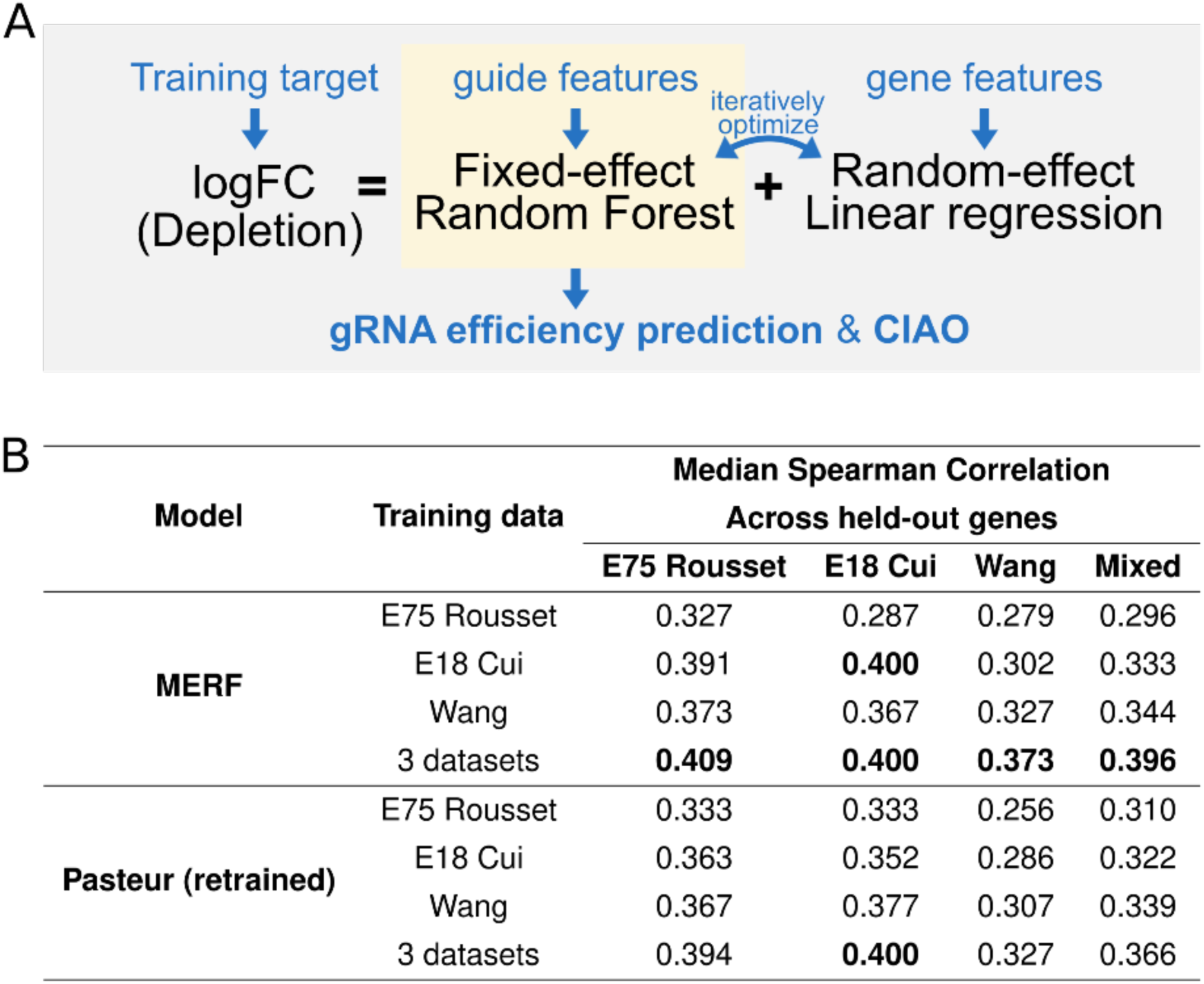
Segregating guide and gene effects produces a predictive model for CRISPRi guide efficiency. (**A**) An overview of the training process of the MERF model. The MERF model segregates depletion values into predictions from a fixed-effect random forest model capturing guide efficiency and a random-effect linear regression model capturing effects associated with the target gene and dataset. The trained fixed-effect random forest model is used for gRNA efficiency prediction and the web-based tool CIAO (ciao.helmholtz-hiri.de). (**B**) Evaluating predictions of guide efficiency after removing gene effects. Spearman correlations between predictions and measured logFC for held-out genes. Genes were held out in 10-fold cross-validation, and the reported median Spearman correlation was calculated across all held-out genes.

Since the fixed effect model from the MERF removes gene-specific effects to estimate guide efficiency, guide-wise cross-validation is not possible, as the true guide efficiency is unknown. As an alternative to guide-wise cross-validation, we developed a gene-wise cross-validation scheme. We trained new models using 10-fold cross validation, this time holding out all guides targeting a set of held-out genes (see also cross-validation, **Methods**; **Table S2**), evaluating the Spearman correlation between predictions and measured depletion within each gene under the assumption that rank order should reflect guide efficiency within a gene. As a comparison, we retrained the previously published Pasteur model [20] on our training sets using the authors’ publicly available source code. Since the Pasteur model was designed to be trained on a single dataset, we implemented a scaling normalization method to make depletion values comparable across datasets (see Data integration, **Methods; Figure S4**) based on prior work on data fusion for Cas9 genome editing in eukaryotes [19]. A comparable normalization factor is calculated automatically within the MERF model through clustering guides targeting the same gene (**Figure S4E**).

As we had previously observed in our evaluation of predictions of guide-wise depletion, data fusion between multiple CRISPRi screens consistently improved performance across models for both the MERF and Pasteur model (**Figure 2B**). The MERF trained on three datasets showed the best overall performance (ρ = 0.396 vs 0.366 for the retrained Pasteur model). We additionally investigated two deep learning models, but neither had improved performance over the MERF (see **Supplementary Note**), possibly due to the limited size of the training data. In sum, the MERF approach provides a straight-forward means of integrating datasets while isolating effects important for guide efficiency.

### Model interpretation with explainable AI illustrates rational design rules for CRISPRi

To understand the features underlying model performance, we again examined SHAP values for our fixed-effect random forest model using TreeExplainer (**Figure 3A**; **Table S8**) [30]. The strongest average effects were seen for distances from the start codon. It has previously been suggested that there is a negative correlation between distance from the start codon and guide efficiency [1], though this has been called into question in subsequent work [31]. To investigate this in our model, we plotted the relationship between the distance to the start codon and SHAP value. The SHAP value plots indicated a strong enhancement of guide activity within 60 bases of the start codon, but only weak effects deeper in the coding sequence (**Figure S5AB&DE**), supporting the absence of a linear relationship between distance and guide efficiency. There was a small but significant difference in the strength of the distance effect in the first 60 bases depending on whether a gene was the first in an operon (**Figure S5C&F**), suggesting that this effect may in part be due to interference with RNA polymerase recruitment or the initial stages of polymerase extension. Accordingly, including an indicator of whether a guide targets the first gene in an operon in our fixed effect model improved cross-validation performance (**Figure S5G**).

**Figure 3:**
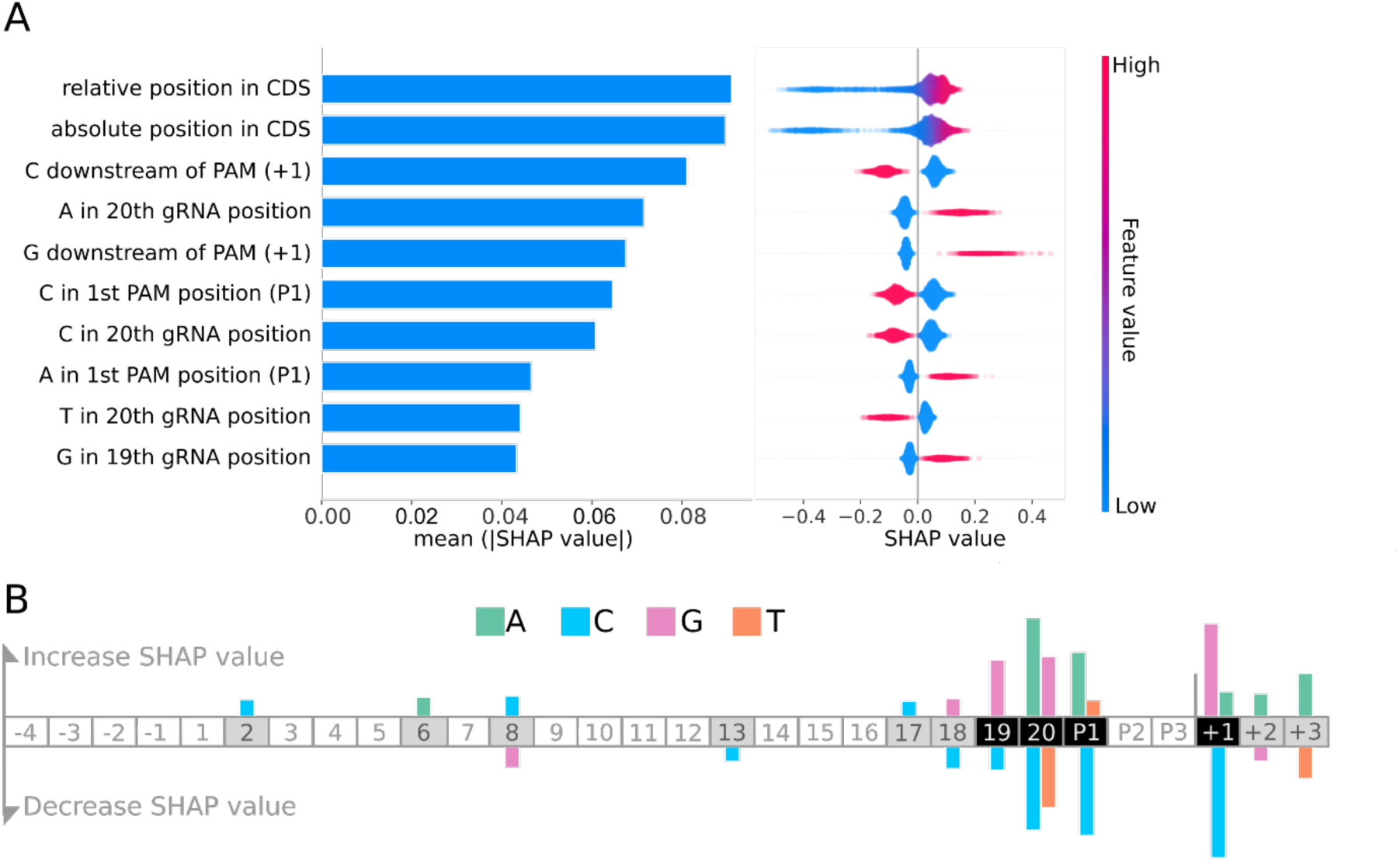
Important features for CRISPRi guide efficiency illustrate sequence preferences. (**A**) SHAP values for the top 10 features from MERF optimized random forest model. Global feature importance is given by the mean absolute SHAP value (left), while the beeswarm plot (right) illustrates feature importance for each guide prediction. CDS: coding sequence.(**B**) A summary of effects of sequence features. Increased SHAP values indicate features that lead to reduced guide efficacy, while decreased SHAP values indicate increased guide efficacy. The guide sequence is numbered G1 to G20 and the three positions of the PAM sequence are labeled P1, P2, P3. Negative and positive numbers refer to positions preceding the guide sequence and following the PAM, respectively.

Other top features involved residues in the vicinity of the PAM sequence (**Figure 3A & B**). For instance, the presence of a cytosine directly downstream of the PAM increased guide efficiency, while guanosine at the same position decreased efficiency as has previously been observed for Cas9-based genome editing in eukaryotes [11]. This may in part be a result of sliding on the PAM sequence leading to formation of disfavored DNA bulges that destabilize the Cas9 complex [33]. Cytosine was favored at the variable position of the NGG PAM, also previously reported for Cas9 editing [11,14]. In contrast, directly before the PAM sequence at position 20 of the guide, guanine and particularly adenine negatively impacted silencing efficiency, while cytosine and thymine increased efficiency — almost the exact inverse of previous reports for Cas9 efficiency in eukaryotic genome editing applications [11,14]. These effects within and around the PAM sequence appeared to interact with each other, as we saw several instances where combinations of features had weaker or stronger effects than would be predicted from the effects of the single features alone (**Table S9**).

### A saturating CRISPRi screen targeting purine biosynthesis genes independently validates performance of tree-based models and data fusion

Our cross-validation results indicated that the MERF-derived fixed-effect random forest trained on multiple datasets outperformed other methods in predicting guide efficiency. As independent validation, we initially explored low-throughput validation approaches, but the resulting data was too sparse to justify strong conclusions (**Figure S6**). To resolve this, we performed a high-throughput screen targeting nine genes from the purine biosynthesis pathway of *E. coli* known to be essential in minimal medium, spread across seven independent transcriptional units (**Figure 4A**). These genes were not included in our initial training set as they are not essential in rich medium, providing a truly independent test set. To avoid any bias in guide selection, we saturated all potential target sites in each gene, ending with a total of 750 gRNAs, including between 35 and 223 guides per gene. Duplicate samples were then collected at three time points during growth in M9 minimal medium, and gRNA depletion was measured with reference to input samples, normalized using a set of 50 gRNAs designed not to target any *E. coli* sequence. Of these nine genes, *purE* and *purK* showed only small variation in guide efficiency (**Figure S7A**), and so were excluded as test genes.

**Figure 4:**
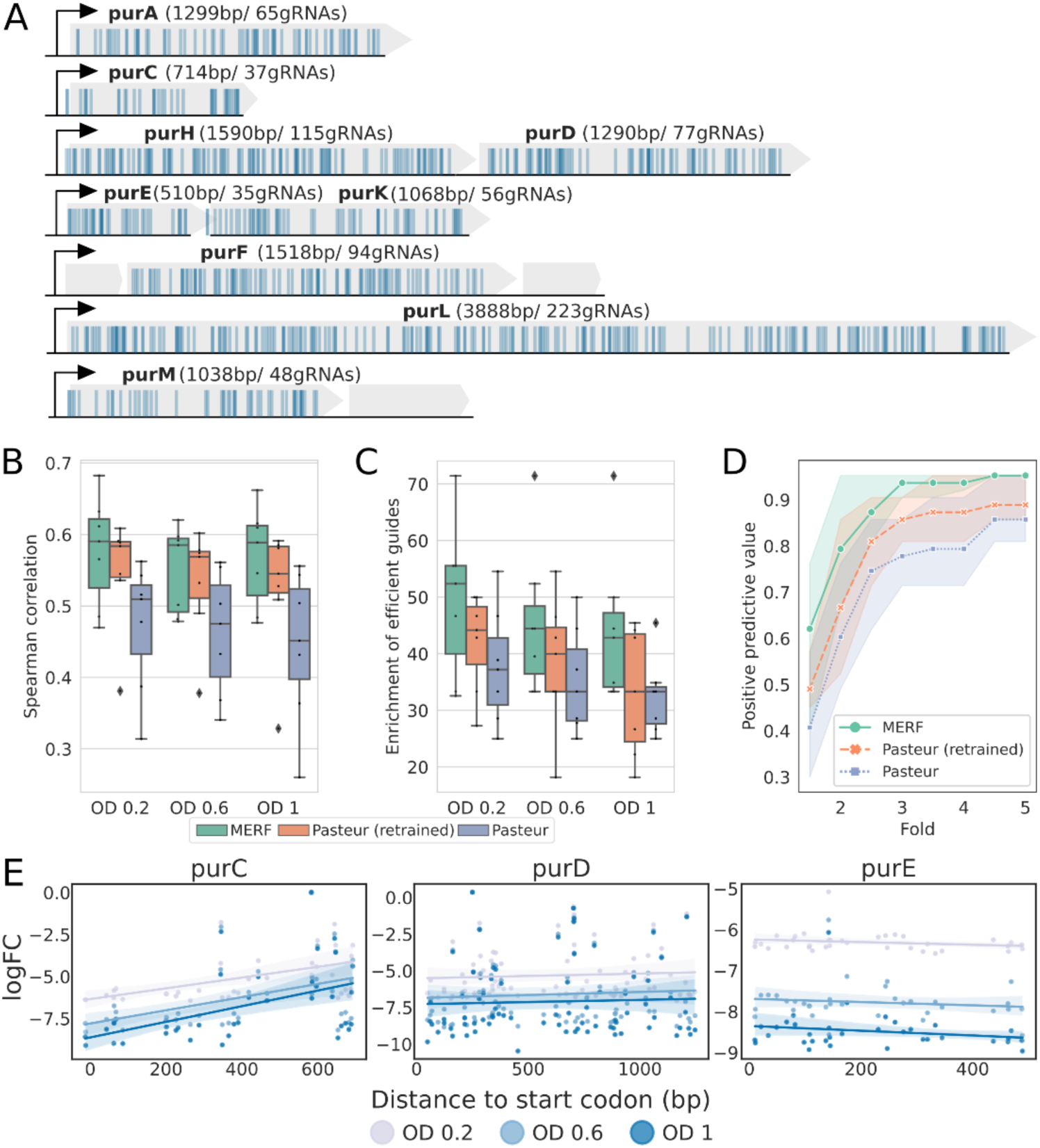
Independent validation of model performance using a saturating screen of purine biosynthesis genes. (**A**) Transcriptional architecture of the 9 targeted purine biosynthesis genes in *E. coli* K12 MG1655. All possible gRNAs were designed for each gene; each blue vertical line represents a gRNA. In total, 750 gRNAs were designed. Grey boxes represent genes, black arrows transcriptional start sites. Spearman correlations between the predicted scores and measured logFC across collected timepoints. **(B) (C)** Enrichment of efficient guides, calculated as the percentage of the experimentally determined 20% most efficient gRNAs in the predicted 20% most efficient gRNAs. The gRNAs are ranked within each gene. (**D**) Positive predictive values for all gRNAs across all time points. The predicted positives are defined as the top 3 predicted most efficient gRNAs for each gene, while the positive class includes gRNAs within N-fold of the depletion value of the most strongly depleted gRNA for each gene (N = 1.5 - 5 with a step size of 0.5). The boundaries of the shaded regions and points indicate PPV values for each time point. The genes *purE* and *purK* were excluded in B-D. (**E**) Measured logFCs for each guide as a function of distance to the start codon for *purC*, *purD*, and *purE*. The shaded regions indicate the 95% confidence interval for the fitted regression line. Plots for the other 6 genes included in the screen are given in **Figure S7B**.

We first compared our MERF-derived fixed-effect random forest to the published Pasteur model and our retrained version of the Pasteur model by calculating Spearman correlation between predictions from each model and the measured logFC for each gene (**Figure 4B**). As in our cross-validation, we saw large boosts in performance for models trained on fused datasets (ρ∼0.478 for the original Pasteur vs. ρ∼0.569 for our retrained Pasteur model), with a smaller further boost in correlation for the fixed-effect model from the MERF (ρ∼0.589). We additionally tested several methods for predicting CRISPRi or CRISPR editing efficiency in eukaryotic organisms, but all exhibited poor performance on our purine screen dataset (**Figure S7D**).

To further investigate differences in performance between the three methods, we devised two additional metrics based on potential use cases. For the first, we calculated how well each method enriches for efficient guides, defined as being in the top 20% of guides for each gene based on observed depletion, mimicking guide selection that might be done in the context of genome-wide screening. Our fixed-effect model led to 10 — 15% increases in median enrichment of efficient guides relative to the original Pasteur model, and 4 — 10% increases relative to our retrained version of the Pasteur model, depending on the selected time point (**Figure 4C**). For the second, we calculated the positive predictive value for each method when selecting the top three guides targeting each gene, mimicking guide selection that might be done when designing guides to selectively silence a particular gene of interest. For the fixed-effect model, there was a 79% chance of selecting a guide that led to depletion within two-fold of the most efficient guide for each gene (**Figure 4D**), compared to 67% and 60% for our retrained and the original Pasteur model, respectively. Given that the most efficient guides we observed across genes were depleted ∼60 — 1,000-fold (**Figure 4E; Figure S7B**), a two-fold deviation still represents substantial depletion. Qualitatively similar differences in PPV between methods were seen using different thresholds for efficiency or when picking the top four or five predicted most efficient guides (**Figure S7C**). In summation, we find that data fusion improves performance across model types in our completely independent validation screen, and that the fixed-effect random forest model from the depletion MERF further increases performance over the state-of-the-art on a variety of realistic tasks.

## Discussion

In this study, we developed a predictive model for CRISPRi guide efficiency using mixed-effect machine learning and integrated data from three gene essentiality screens in *E. coli*. Our approach was based on two primary insights: that effects from the targeted gene dominate over the effects of guide efficiency in depletion screens, and that fusing data across multiple screens can improve prediction accuracy and generalizability. We trained a mixed-effect random forest model (MERF) that uses a linear random-effect model to capture the confounding effects of the targeted gene and dataset, and a fixed-effect random forest that captures the residual effect of guide efficiency on depletion. We showed that this fixed-effect random forest model improves on the previous state-of-the-art using both gene-wise cross-validation on our training data as well as a fully independent screen of guides targeting purine biosynthesis genes essential in minimal medium. These investigations provide a blueprint for developing similar predictive models in the absence of direct measurements of guide efficiency, both for other CRISPR-Cas systems [34] and technologies, as well as for CRISPRi in different bacteria where design rules may vary. We have made a web server for predicting CRISPRi guide efficiency using our MERF publicly available at: https://ciao.helmholtz-hiri.de.

Beyond developing a predictor for CRISPRi guide efficiency, our process of model development and validation provided several new insights into the behavior of CRISPRi screens. Strikingly, we found that gene expression was the single largest contributor to predictions of gene depletion as measured by SHAP values, where higher expression was associated with higher depletion. As the availability of transcriptomics data may be lacking for some organisms, we also tested the possibility of using the codon adaptive index (CAI) as a proxy, with promising results (**Figure S7 EF&G**), indicating that this observation is not an artifact of the expression dataset we used.

Similarly, our saturating purine validation screen revealed previously unobserved features of CRISPRi depletion screens. It has been suggested that CRISPRi guide efficiency is highest near the transcription start site and declines further into the coding sequence [1,25]. In contrast to this, we saw at least three distinct patterns of depletion across genes (**Figure 4E & S7B**): *purC* and *purM* showed the expected trend towards decreasing efficiency along the gene; the majority of genes had good and bad guides distributed across the length of the gene with no clear positional preference; and *purE* and *purK* showed very little variation in guide efficiency. These differences in the relationship between distance to the start codon and guide efficiency were surprising, as distance features were clearly important to our model predictions (**Figure 3A**). We attempted to train a model excluding distance features, but this substantially degraded performance on predicting depletion in our high-throughput screen (**Figure S7EG&F**). Investigation of the effects of our distance feature on predictions indicate a sensitive region of ∼60 bases from the start codon where silencing is more effective, possibly associated with transcription start sites or the early steps of transcriptional elongation (**Figure S5**); additional work will be required to confirm this. In sum, our screen of guides targeting purine biosynthesis genes highlighted some unexpected features of CRISPRi while also independently validating the improved performance of our random forest model compared to the state-of-the-art.

Prediction of guide efficiency will become increasingly important with more complex applications of CRISPR technologies. In particular, the potential for multiplexing CRISPRi could be transformative when compared to established technologies. One example of this would be in screening for fitness interactions between genes. The current state-of-the-art is based on arrayed mating of single gene deletion libraries [35,36], which is both labor intensive and technically challenging, and becomes increasingly so when querying higher-order interactions [37]. A similar example is in metabolic engineering where multiplexed CRISPRi can be used to modulate biosynthetic pathways to optimize production of a particular metabolite for industrial applications [38]. The development of CRISPR array technologies that can co-express as many as 22 guides simultaneously [9,10] should accelerate the development of these approaches. However, large-scale, multiplex applications will require better tools for guide design to ensure robust results. Individually screening guides for activity quickly becomes prohibitive when one considers applications that require hundreds or thousands of guides. The machine learning approach presented here provides a straight-forward solution to this problem.

While we focused here on applications of CRISPRi with dCas9 in *E. coli*, the techniques we have developed are in principle generic and could be extended to CRISPRi with any catalytically-dead nuclease in any bacterium of interest, or even to entirely different CRISPR technologies. For instance, we recently applied the same basic methodology to investigate the features underlying autoimmune activation of Cas13 targeting cellular RNA [34]. It is becoming increasingly clear that the performance of CRISPRi depends on both genetic background and the specific Cas protein used. For instance, *Streptococcus pyogenes* dCas9 expression has low silencing efficiency in some bacteria and can even be toxic [39], forcing the adoption of alternative Cas effector nucleases [40]. Alternative Cas effectors have large differences in their PAM preferences and the stringency of the PAM requirement [41]; presumably, alternative dCas proteins may also respond differently to the other gene and guide features described here. The approach outlined here, applying autoML and explainable AI to rapidly arrive at a description of the design rules underlying the efficiency of CRISPRi silencing, provides a means to rapidly characterize the behavior of new dCas proteins as genome-wide screening data becomes available.

## Supplementary Figures

**Figure S1:**
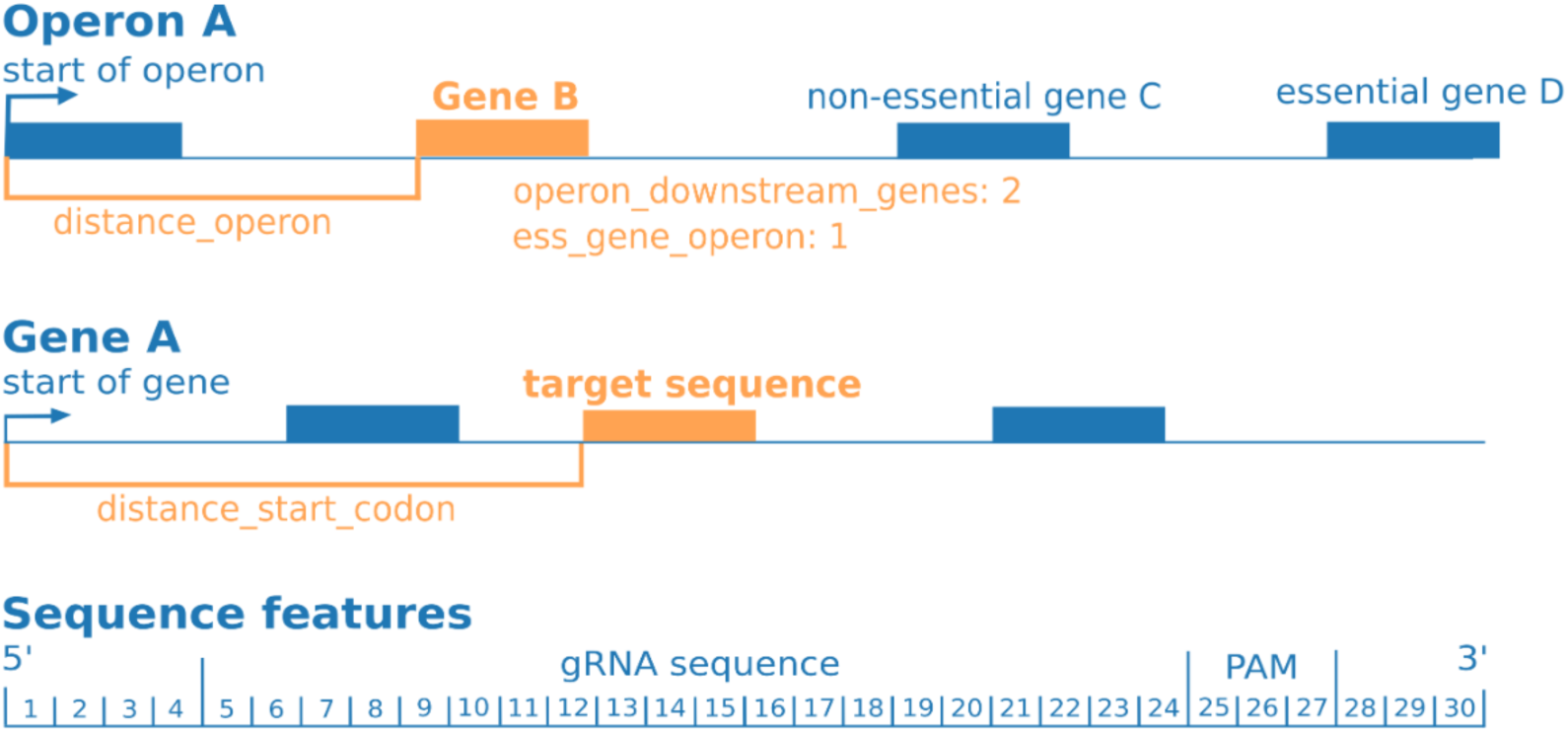
Illustration of the genomic and sequence features used, see also Table S1.

**Figure S2:**
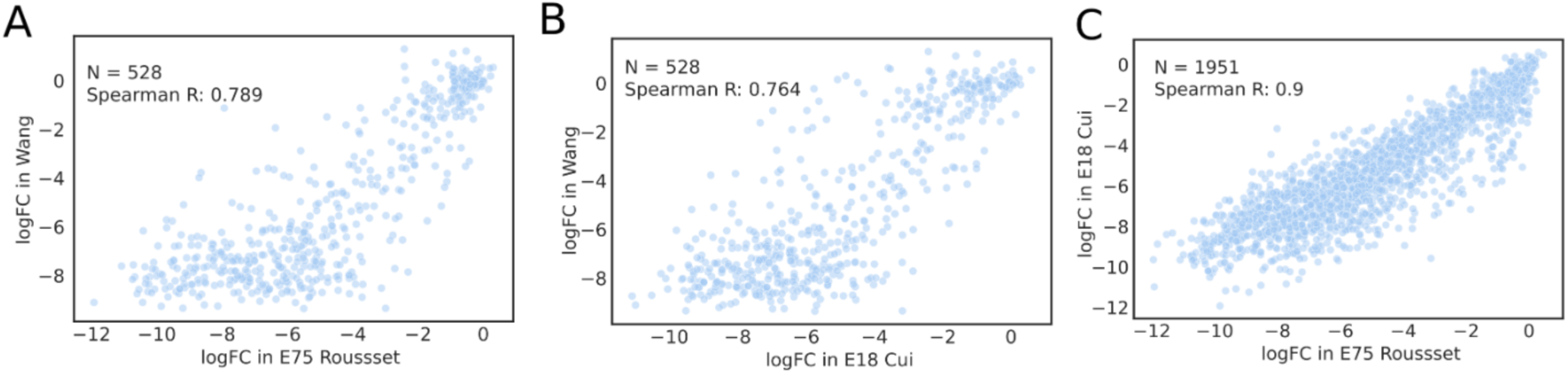
Comparison of guide depletion across datasets. (**A**) The logFC of gRNAs in E75 Rousset plotted against that in Wang for shared gRNAs. (**B**) The logFC of gRNAs in E18 Cui was plotted against that in Wang for shared gRNAs. (**C**) The logFC of gRNAs in E18 Cui was plotted against that in E75 Rousset for overlapping gRNAs.

**Figure S3:**
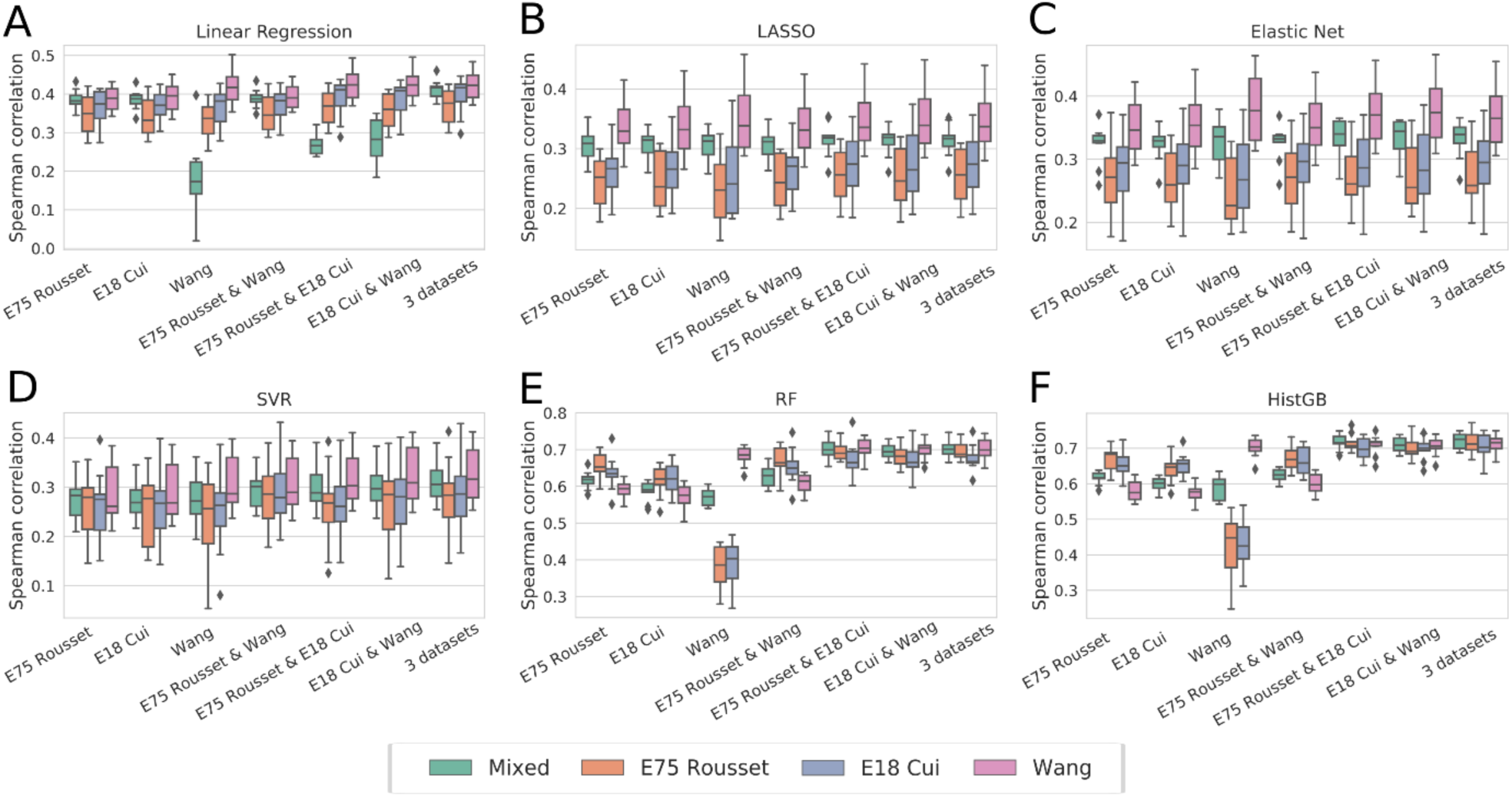
Spearman correlation of 10-fold cross-validation of models trained with one or mixed datasets. (A) linear regression, (B) LASSO, (C) Elastic net, (D) support vector regression (SVR), (E) Random forest (RF) regression, (F) Histogram-based gradient boosting regression.

**Figure S4:**
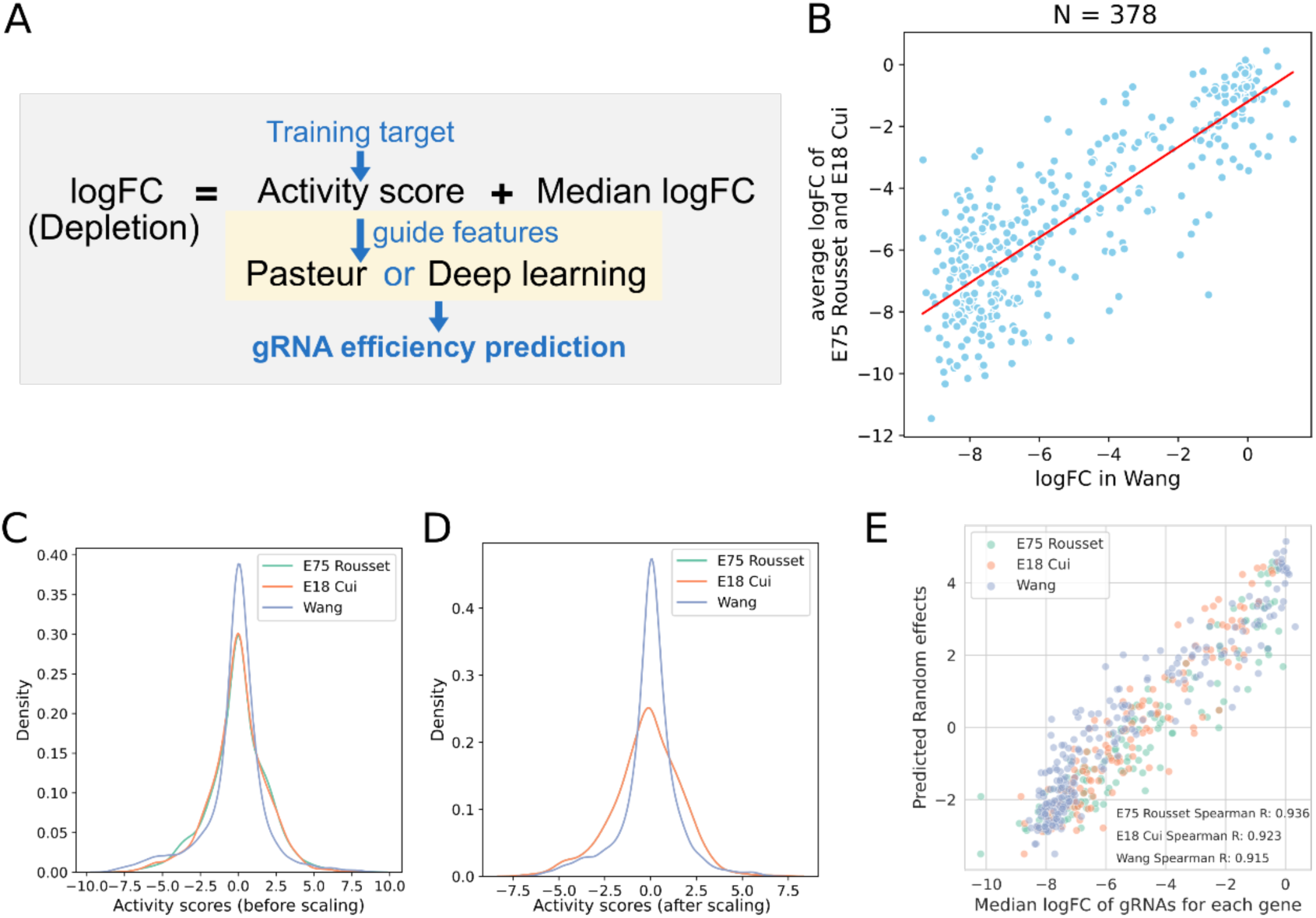
Data integration for retrained Pasteur and deep learning models. (**A**) The overview of retraining the Pasteur and deep learning models. We subtract the gene-wise median logFC from each gRNA depletion value upon data fusion to obtain the activity scores of each gRNA. The activity scores were used as training targets and the Pasteur model and deep learning models were trained with guide-specific features. (**B**) The logFC values in Wang were scaled based on a linear regression between the original logFC of Wang and the average logFC of E75 Rousset and E18 Cui for 378 overlapping gRNAs. The distributions of activity scores (**C**) with and (**D**) without scaling are shown. (**E**) Predicted scores of the random effect model from MERF (y-axis) compared to the median logFC across gRNAs (x-axis) for each gene in each dataset.

**Figure S5:**
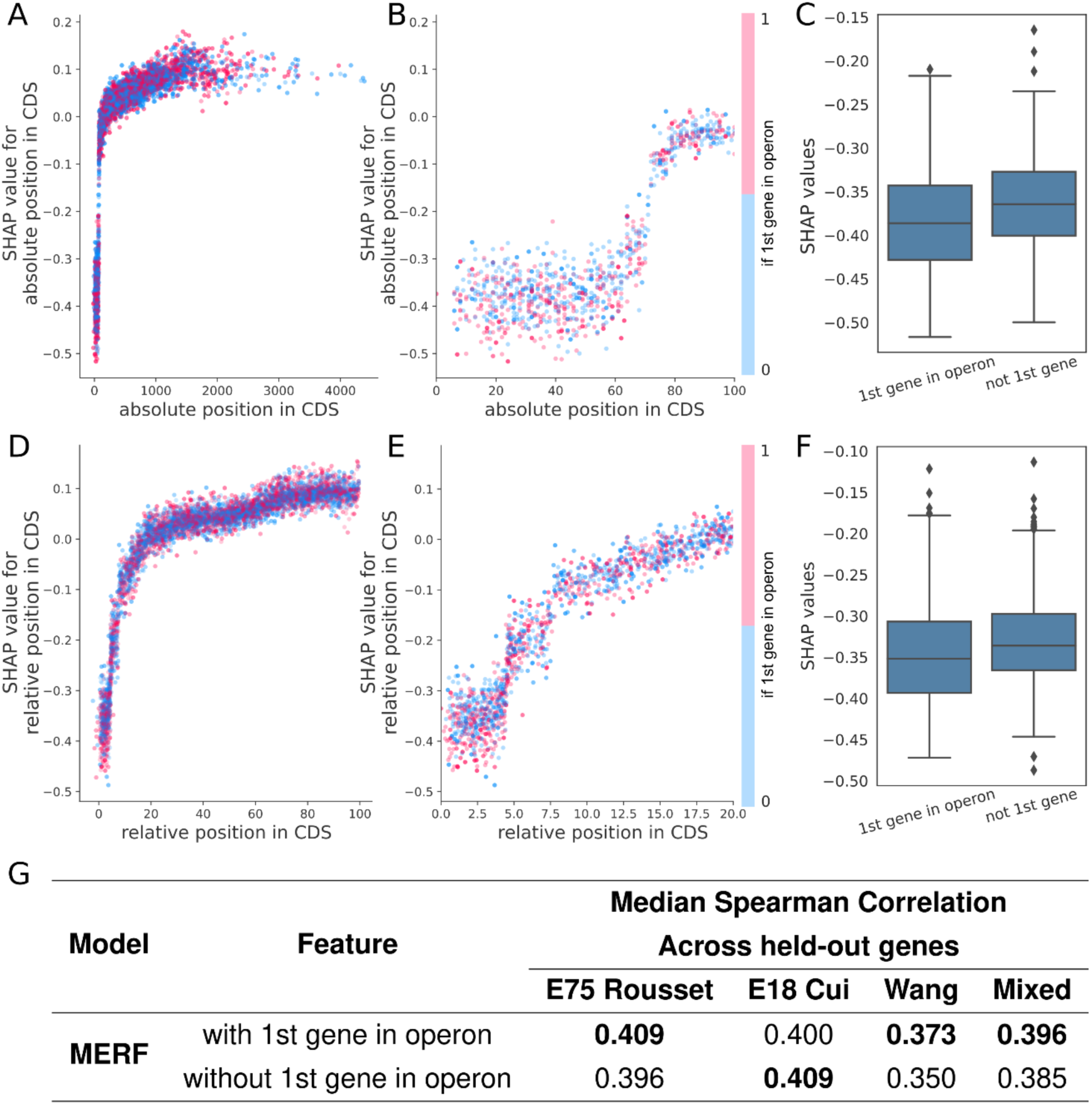
Interaction between distance features and whether targeting gene is the first gene in operon. (**A-B**) SHAP values for absolute position in CDS plotted against absolute position. The dots are colored based on whether the guide is targeting the 1st gene in operon. Red indicates the guide targets the 1st gene in operon, while blue indicates it is not. In **B**, the x-axis is limited to range 0-100. (**C**) SHAP values for absolute position in CDS of guides targeting within the first 70 bases. SHAP values are significantly lower for 1st genes in operons (p = 2.13 X10^-8^, two-sided Mann-Whitney U test). (**D-E**) Similar to A-B but for relative position in CDS. In **E**, the x-axis is limited to range 0-20. (**F**) SHAP values for relative position in CDS of guides targeting relative position in CDS lower than 5%. SHAP values are significantly lower for 1st genes in operons (p = 1.68 X10^-5^, two-sided Mann-Whitney U test). (**G**) Evaluating predictions of guide efficiency after removing gene effects for MERF models trained with or without the feature of whether the target gene is the 1st gene in operon, using the same cross-validation procedure as Figure 2B.

**Figure S6:**
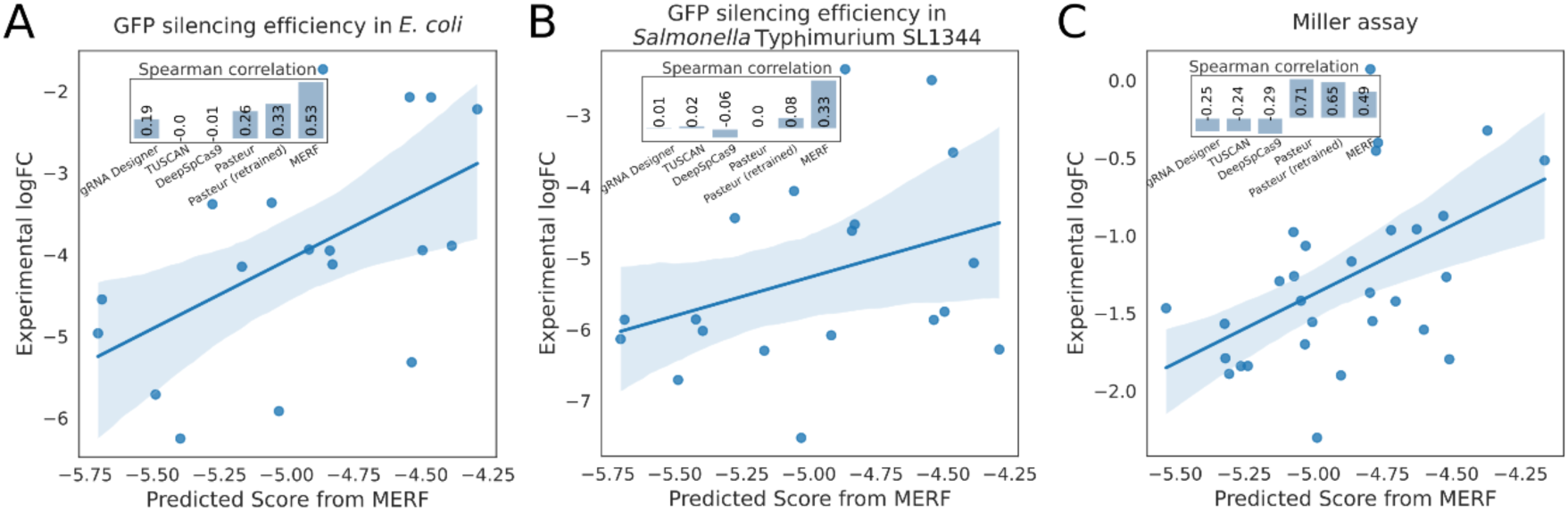
Independent low-throughput validation of model performance. The activity of 19 gRNAs targeting a plasmid-expressed deGFP gene was measured in (**A**) *E. coli* and in (**B**) *Salmonella* Typhimurium SL1344 using a flow cytometry-based assay. The measured activity compared to the control gRNA is plotted against the score predicted by the MERF model. The inset barplot illustrates Spearman correlations for six methods for predicting guide efficiency. (**C**) The activity of 30 gRNAs targeting lacZ was measured with a Miller assay by [20], plotted as in A and B.

**Figure S7:**
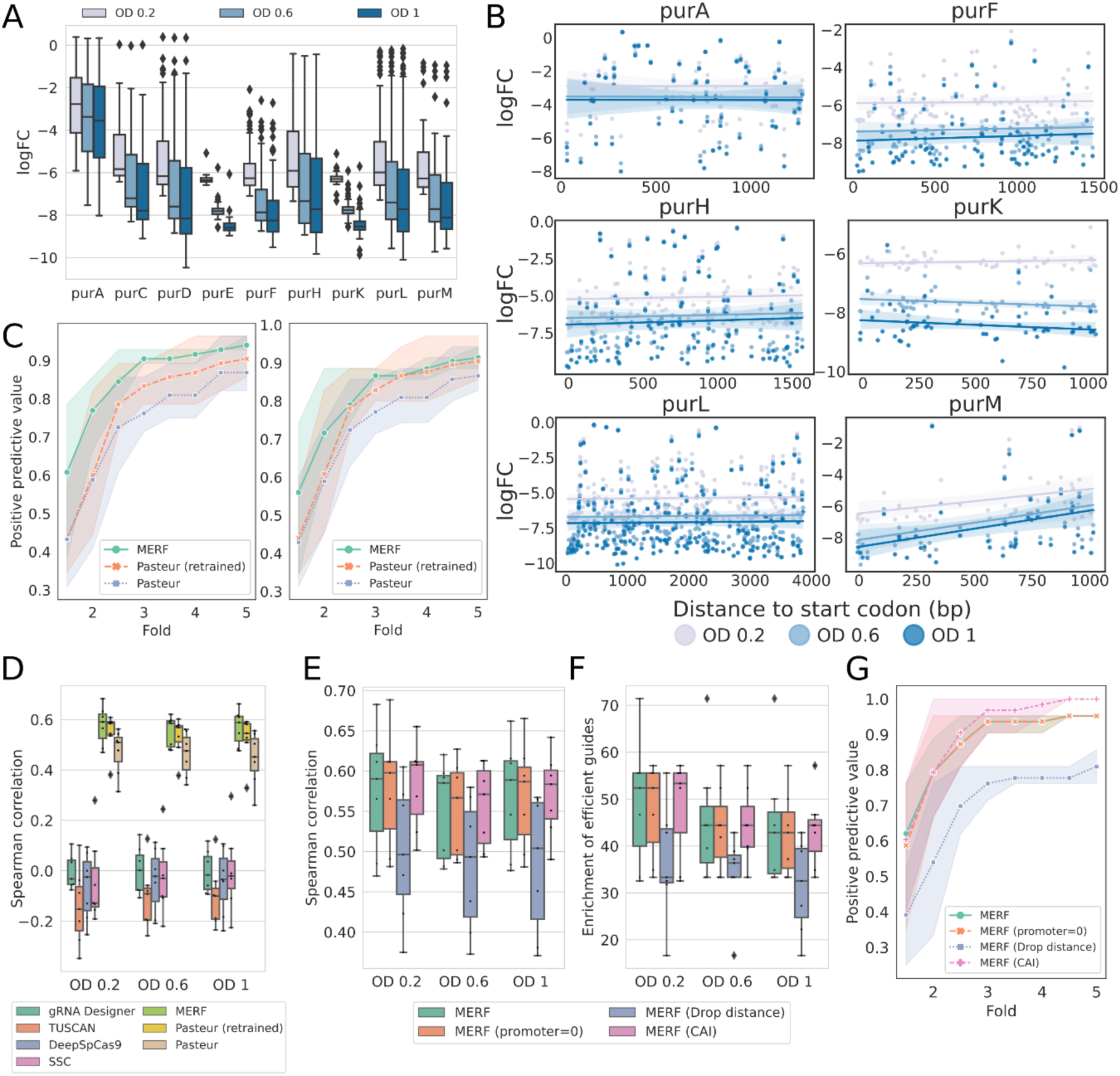
Additional figures related to the saturating screen of purine biosynthesis genes. (**A**) The distribution of experimental logFC at different time points for each gene. (**B**) Measured logFCs for each guide as a function of distance to the start codon for the other 6 genes not shown in Figure 4E. (**C**) Positive predictive values of all gRNAs for each time point. The predicted positives are defined as (**left**) the top four and (**right**) the top five predicted gRNAs in each gene, while true positives are gRNAs with fold change within the N fold of the strongest depleted gRNA in each gene (N= 1.5 - 5 with a step of 0.5). (**D**) Spearman correlations between the predicted scores and measured logFC across collected timepoints. (**E-G**) Performance on the purine screen of MERF models trained with the promoter feature set to 0 (promoter=0), without distance features (Drop distance), and with 4 instead of 9 gene features for the random-effect model (CAI value, gene length, gene GC content, and dataset). The calculation of Spearman correlation, enrichment of efficient guides, and positive predictive value are the same as Figure 4B-D.

**Figure S8:**
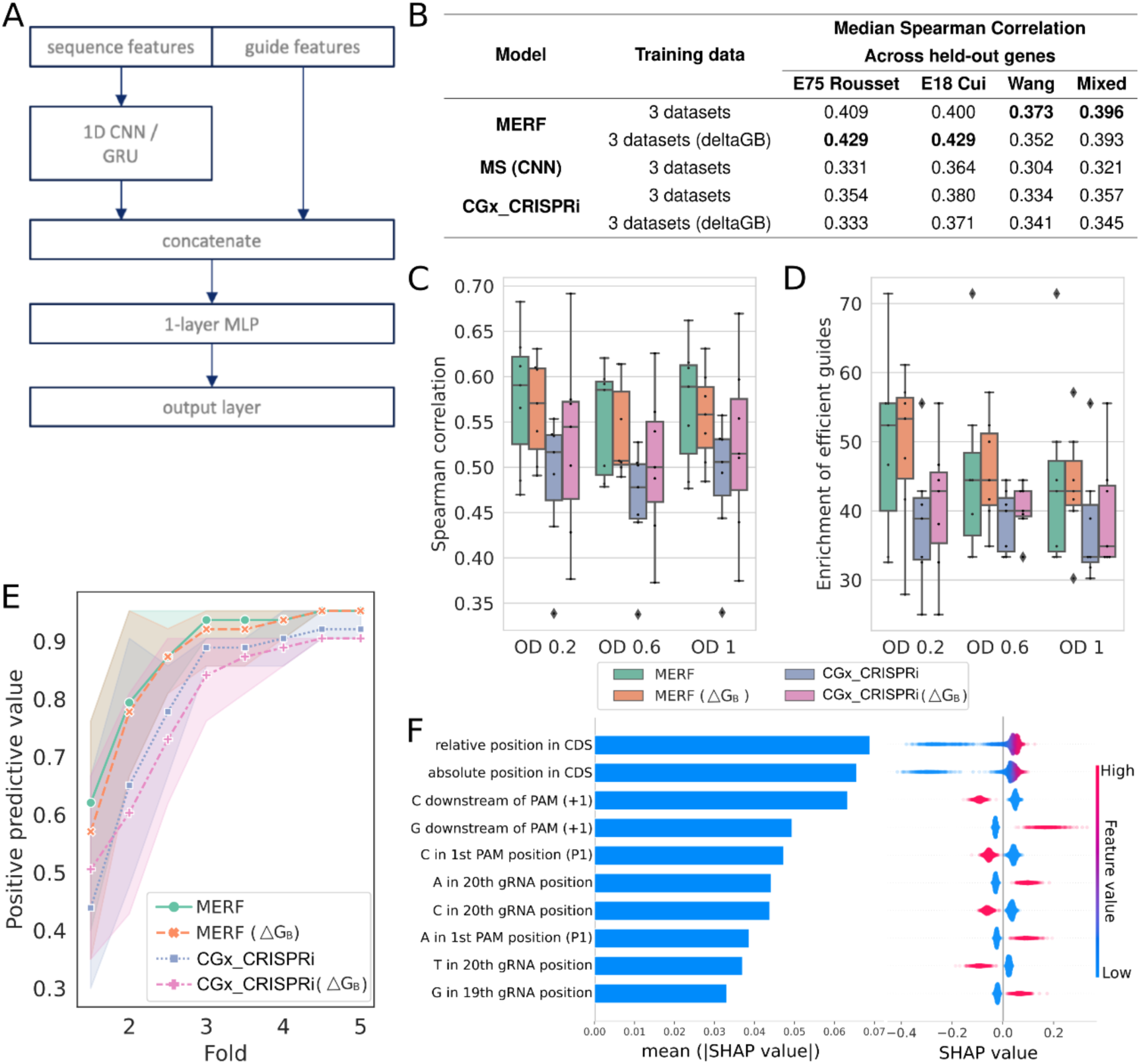
(**A**) Architectures of the applied deep learning models. Guide features refer to guide-specific features apart from sequence features. MLP: multilayer perceptron. (**B**) Evaluating predictions of guide efficiency after removing gene effects. Spearman correlations between predictions and measured logFC for held-out genes. Genes were held out in 10-fold cross-validation, and the reported median Spearman correlation was calculated across all held-out genes. When △G_B_ is specified, the MERF or CGx_CRISPRi models were trained with △G_B_ instead of the original four thermodynamic features. (**C-E**) Performance on the purine screen of MERF and CGx_CRISPRi models. When △G_B_ is specified, the MERF or CGx_CRISPRi models were trained with △G_B_ instead of the original four thermodynamic features. The calculation of Spearman correlation, enrichment of efficient guides, and positive predictive value are the same as Figure 4B-D. (**F**) SHAP values for the top 10 features from MERF model trained with △G_B_. Global feature importance is given by the mean absolute SHAP value (left), while the beeswarm plot (right) illustrates feature importance for each guide prediction.

## Supplementary Tables

Table S1 Description of the features used in model training Table S2 Description of train-test splits in the study

Table S3 Model evaluation of feature engineering

Table S4 SHAP values from model predicting the gRNA depletion Table S5 Model evaluation of data fusion

Table S6 Model evaluation of data fusion with other model types Table S7 Model evaluation of segregating gene and guide effects Table S8 SHAP values from segregation models

Table S9 SHAP interaction values from MERF Table S10 Description of gRNAs targeting deGFP Table S11 Description of gRNAs in miller assay

Table S12 Model evaluation for targeting deGFP and Miller assay

Table S13 Description of gRNAs for screen of purine biosynthesis genes Table S14 Model evaluation of saturating screen of purine biosynthesis genes Table S15 List of strains

Table S16 List of plasmids

Table S17 List of oligos for plasmid construction, library generation, and sequencing

## Supplementary Note: Deep learning approaches do not improve prediction performance

Given that we saw better performance with tree-based methods over linear regression, we asked if model complexity was a limiting factor in prediction. Deep learning approaches have been applied to predicting guide efficiency for a number of CRISPR technologies [16–19,42]. Considering this, we asked if deep learning models would also improve performance in predicting gRNA efficiency for CRISPRi in bacteria. As a representative architecture, we implemented a one-dimensional convolutional neural network (CNN), which runs a series of kernel filters across the sequence to extract local features. As a second model representative of current deep learning approaches, we reimplemented a state-of-the-art deep learning architecture based on the CGx architecture used by CRISPRon [19] for Cas9 genome editing in eukaryotes, only trained using our CRISPRi data and using our feature set. Both models were trained using scaled data as done for re-training the Pasteur model (see Data integration for retrained Pasteur and deep learning models, **Methods**).

For our custom CNN architecture, we used the convolutional layers to extract sequence features before concatenating them to the rest of our guide feature set (**Figure S8A**). This concatenated feature set was then fed through a fully connected 4-layer multilayer perceptron (MLP) for regression using MS values for guide efficiency. Both the custom CNN and CGx_CRISPRi models exhibited lower Spearman correlations as compared to our previously trained random forest models when tested on held-out gene sets (**Figure S8B**; Table S7; CNN ρ=0.321, CGx_CRISPRi ρ=0.357, vs. MERF ρ=0.396). As one major difference between our CGx_CRISPRi implementation and that of CRISPRon is in the use of the △G_b_ parameter that integrates a complete thermodynamic model of Cas9 binding [43], we additionally tested substituting this for our simpler thermodynamic features in both CGx_CRIPSRi and our MERF model, leading to no clear improvement in prediction performance (**Figure S8BCD&E**). These results show that conventional machine learning approaches can outperform deep learning architectures and suggest that data may currently be limiting for more complex machine learning approaches, though it remains possible that specialized architectures may improve on the approaches implemented here.

## Methods

### Training datasets

We collected the data from three previous CRISPRi genome-wide essentiality screens in *E.coli* K12 MG1655 [21,25,31]. The sequence, targeted gene, gene position, and fitness effect of each gRNA were retrieved from the supplementary information of each study. Gene sequences and positions were updated to be consistent with the latest reference genome version (NCBI: NC_000913.3). We discarded gRNAs from the Wang data set previously removed as having insufficient read counts [25] or sequences from the Rousset and Cui datasets that differed from the reference sequence due to differences in the genome versions. 8099 gRNAs targeting the coding-strand within the coding regions of essential genes were extracted in total from all three datasets.

### Feature engineering

A Python script (feature_engineering.py) was used to compute 137 sequence, thermodynamic, genomic, and transcriptomic features (**Table S1**). 30-mer sequences, from 4 bp upstream of gRNA to 3 bp downstream of the PAM sequence, were one-hot encoded to 120 (30 × 4) features. Thermodynamic features including minimum free energy for different interactions (the hybridization of gRNA and target DNA, the hybridization of the seed region of gRNA and target DNA, the homodimer of gRNA, and the monomer of gRNA) were computed using the ViennaRNA Package [26]: RNAduplex (version 2.4.12) for RNA:RNA hybrids; RNAduplex (version 2.1.9h) for DNA:RNA hybrids [27]; RNAfold (version 2.4.12) for single RNA folding. The seed region was defined as the 8 nt PAM-proximal region in the gRNA. The CRISPRoff score (deltaGB) was calculated using the energy function in the CRISPRoff pipeline v1.1.2 [43] with ViennaRNA Package version 2.2.5 (deltaGB_calculation.py). Homopolymer was defined as the number of consecutive repetitive nucleotides in the gRNA sequences. Genomic features including gene and operon organizations were based on the reference genome, essential genes as determined in the Keio collection [24], and transcriptional unit definitions from RegulonDB [44]. Transcriptomic data including gene expression levels across growth at ten different ODs were obtained from a previous study [28]. Minimal or maximal expression levels were calculated across the range of ODs until the growth phase when cells were collected in each CRISPRi screen: OD 1.4 for the Wang dataset, and all ODs for the Rousset and Cui datasets. The codon adaptation index (CAI) for each gene was calculated using CAIcal [45]. The resulting feature sets are available on GitHub.

### Cross-validation for machine learning methods

To evaluate the models for depletion prediction, training and test sets were split guide-wise based on unique gRNA sequences. 10-fold cross-validation was used to evaluate model performance. 10 test sets, with the number of samples ranging from 786 to 855 targeting 245 to 262 genes, were kept identical regardless of training data. The minimum and median Hamming distance were 6 and 21 between the train and test sets in each iteration and between test sets, as calculated with the hamming function from SciPy (version 1.10.0) (Virtanen et al., 2020). For each iteration, the Spearman correlation between measured depletion values and predicted values for all test samples was calculated.

To evaluate the models for gene efficiency prediction, training and test sets were split by gene based on gene identifier. 10-fold cross-validation was used to evaluate model performance. 10 test sets, with the number of samples ranging from 624 to 806 gRNAs targeting 30 or 31 genes, were kept identical regardless of training data. The minimum and median Hamming distance between the train and test sets in each iteration and between test sets were 7 and 21. For each iteration, the Spearman correlation between measured depletion values and predicted values for each held-out gene was calculated. For MERF, predictions were performed using the fixed-effect model. The gRNAs used in each train-test split are listed in **Table S2**.

### Predictive models for depletion

The automated machine learning toolkit auto-sklearn (version 0.10.0) [22] was used to develop optimized machine learning regression models. For auto-sklearn, the AutoSklearnRegressor function was used and all possible estimators were included. The following parameters were used: “time_left_for_this_task” = 3600, “per_run_time_limit” = 360, “resampling_strategy” = ‘cv’, ensemble_kwargs = {“ensemble_size”: 1}, “resampling_strategy_arguments” = {“fold”: 10}, “metric” = autosklearn.metrics.mean_squared_error. Feature types for each feature are listed in **Table S1**. The selected models were saved and used with scikit-learn for downstream analysis.

Simple linear regression, LASSO, elastic net, SVR, random forest, and histogram-based gradient boosting models were trained using scikit-learn (version 0.22.2) [23].

### Segregation of guide and gene effects with MERF

We removed genes with less than 5 gRNAs in each dataset to stabilize estimates of median gRNA activity scores for a fair comparison between the MERF and models requiring scaling data integration (see below), resulting in 7400 gRNAs in total. This included 1618 gRNAs targeting 171 genes from E75 Rousset/E18 Cui and 4164 targeting 300 genes from Wang.

MERF models were trained using the package merf (version 1.0). Hyperparameters for the final fixed-effect random forest model were optimized using hyperopt (version 0.2.5) [46]. The search space included: ‘bootstrap’ either True or False, ‘n_estimators’ from 50 to 1000 with a step of 10, ‘max_features’ from 0 to 1, ‘max_depth’ from 2 to 30 with a step of 1, ‘min_samples_leaf’ from 1 to 20 with a step of 1, ‘min_samples_split’ from 2 to 20 with a step of 1 (for more details, see MERF_crispri.py on GitHub). 129 guide-specific features were assigned as fixed effects, while 9 gene-specific features were assigned as random effects. The random effect matrix was standardized with the default StandardScaler function from scikit-learn. 301 unique gene IDs were used as cluster IDs. To train simplified models excluding transcriptomic measurements (**Figure S7E-G**), the CAI value, gene length, gene GC content, and dataset were included for the random-effect model. The output from the complete MERF model was in the range of -12 to +2, and output from the fixed-effect random forest was in the range of -7 to -2.5.

In all cases, the fixed effect was used for prediction (i.e. in cross-validation and validation with the purine screen), requiring only features associated with the guide sequence.

### Data integration for retrained Pasteur and deep learning models

For the Pasteur and deep learning models, logFC values were scaled to integrate the datasets, adapted from a previously applied data fusion method [19]. First, the mean of logFCs of E75 Rousset and E18 Cui were calculated and used as the scaled logFC (**Figure S4B-D**). Then linear regression was performed between the logFCs in Wang and scaled logFCs in E75 Rousset for 378 overlapping gRNAs. All of the logFCs from Wang were then scaled by the fitted slope and intercept values (**Figure S4B**). The 378 overlapping gRNAs in Wang were excluded in the subsequent training. Activity scores were calculated by subtracting the scaled logFC of each gRNA from the median scaled logFC for each gene across all 3 datasets (**Figure S4A**). For cross-validation, scaling was performed within each test fold to avoid possible leakage of information between test and training sets.

### Deep learning model implementation and training

Deep learning models were trained using pytorch (version 1.8.1) [47] and pytorch-lightning (version 1.5.10). For our custom 1D CNN model, sequence features were processed using 1D convolutional layers and later concatenated with other guide features (**Figure S8A**). Concatenated features were further processed with fully connected layers. Three 1D convolutional layers were implemented sequentially with input channels 4, 64, and 64, output channels 64, 64, and 32, kernel size 5, 3, and 1, and stride 2, 2, and 1 respectively. For fully connected layers, output dimensions are 128, 64, 32, and 1 (which is predicted gRNA efficiency). The first three fully connected layers are accompanied by batch normalization [48], ReLU, and dropout [49] (p=0.5). We trained the model using AdamW [50] optimizer with learning rate of 0.001 and batch size of 32.

For CGx_CRISPRi, we implemented the CRISPRon architecture (CGx) modified to include our nine non-sequential guide features (distance features, thermodynamic features, etc.) concatenated to the processed sequential features. To test the effect of incorporating the deltaGB score [43], the four thermodynamic features were replaced with deltaGB, resulting in concatenating five non-sequential guide features to the processed sequential features. We trained the model using the Adam optimizer [51] with a learning rate of 0.001 and batch size of 32. We additionally tested a learning rate of 0.0001 and batch size of 500 as used in the original CRISPRon implementation [19], but saw only minor differences in performance (Table S7).

### Model interpretation

Tree-based models, including depletion prediction models, and the fixed-effect model from MERF were interpreted using TreeExplainer from the python shap package (version 0.39.0) [30].

SHAP values were calculated using the ‘shap_values’ function in TreeExplainer with 80% of the samples for depletion prediction models and all samples for guide efficiency. SHAP value plots were generated with the ‘summary_plot’ function in shap.

SHAP interaction values were calculated using the ‘shap_interaction_values’ function in TreeExplainer with 1000 guides. Absolute SHAP interaction values were averaged over 1000 samples. The rank of interaction was obtained based on the sorted mean absolute SHAP interaction values across all unique feature pairs. To compare interaction effects to expectations based on single-feature SHAP values, four feature combinations were considered: both absent (-/-), only the first feature present (+/-), only the second feature present (-/+), and both present (+/+). For the top 5,000 interacting feature pairs, the SHAP values for each feature in samples with each combination of features were extracted. For each feature pair (F1 and F2), the expected value for +/+ was calculated as the sum of the median F1 SHAP values for +/- samples with the median of F2 SHAP values for -/+ samples, while the expected value for -/- was calculated as the sum of the median F1 SHAP values for -/+ samples and the median of F2 SHAP values for +/- samples.

### Strains and growth conditions

All strains, plasmids, and primers are listed in Supplementary **Table S15, S16, and S17**. *E. coli* cells were grown in Lysogeny Broth (LB) (10 g/L NaCl, 5 g/L yeast extract, 10 g/L tryptone) at 37 °C with shaking at 250 rpm. To maintain plasmids, the antibiotics ampicillin, chloramphenicol, and/or kanamycin were added at 50 µg/mL, 34 µg/mL, and 50 µg/mL, respectively as necessary. For screening experiments, *E. coli* MG1655 was grown in M9 minimal medium (1x M9 salts, 1 mM thiamine hydrochloride, 0.4% glucose, 0.2% casamino acids, 2 mM MgSO4, 0.1 mM CaCl2) supplemented with the appropriate antibiotics.

### Validation of GFP silencing by flow cytometry in *E. coli* and *S.* Typhimurium

To investigate gene repression efficiency, 19 sgRNAs were selected to target the coding strand of a *degfp* reporter gene at different positions in *E. coli* BL21(DE3) (**Table S15**). Cells were initially transformed with three compatible plasmids encoding dCas9, a *degfp*-targeting sgRNA, and a deGFP reporter (**Table S17**). For normalization purposes, a positive control strain harboring a non-targeting sgRNA and a negative control strain lacking the *degfp* encoding reporter plasmid was included. Overnight cultures of cells harboring the above-mentioned plasmids were back-diluted to OD_600_ ∼0.01 in LB medium with ampicillin, chloramphenicol and/or kanamycin and incubated with shaking at 250 rpm at 37 °C, until reaching an OD_600_ of 1. Cultures were then diluted 1:25 in 1x phosphate-buffered saline (PBS) and analyzed on an Accuri C6 flow cytometer with C6 sampler plate loader (Becton Dickinson) equipped with CFlow plate sampler, a 488-nm laser, and a 530+/- 15-nm bandpass filter. Forward scatter (cutoff of 11,500) and side scatter (cuttoff of 600) were used to eliminate non-cellular events. The mean green fluorescence value (measured by the FL1-H channel) across 30,000 events within a gate set for *E. coli* was used for further analysis. The log fold repression of each gRNA was calculated as the ratio between the difference in fluorescence values between the gRNA and negative control, and the difference between the positive and the negative control, followed by log transformation. The mean log fold repression across three replicates was compared to predicted values from the machine learning models (**Table S10 and S12**).

For experiments with *S.* Typhimurium, the procedure was similar, but cells were grown until an OD_600_ of ∼0.8 before analysis on an Accuri C6 flow cytometer. To eliminate non-cellular events, the forward scatter (cutoff of 10,000) was used and the mean green fluorescence value (FL1-H) across 30,000 events within a gate set for *S.* Typhimurium was used for data analysis as described above across four replicates (**Table S10 and S12**).

### Generation of the sgRNA library for purine biosynthesis genes

9 genes (*purA*, *purC*, *purD*, *purE*, *purF*, *purH*, *purK*, *purL*, *purM*) in the purine biosynthesis pathway in *E. coli* MG1655 were selected for the saturating screen in M9 minimal medium. All possible 20 nt gRNAs with an NGG PAM, GC content between 30% and 85%, and without BbsI restriction sites were included in the library, resulting in 750 gRNAs (**Table S13**). The minimum and median Hamming distance of the 30-mer sequences between the 750 gRNAs and the training data from three essentiality screens were 7 and 21.

For the sgRNA library, plasmid DC512 served as a backbone, following a previously established protocol [9]. To generate a library with 800 sgRNAs (including 50 randomized non-targeting sgRNAs; **Table S13**), 800 forward and reverse oligonucleotides each encoding one spacer and a 4-nt junction, were synthesized as an oPool (1600 oligos at 10pmol/oligo) by Integrated Device Technology (IDT). The same 5′ and 3′ assembly junction sequences were used for all spacer pairs leading to the same integration site within the backbone (5′ TAGT overhang at the 5′ end and a 3′ AAAC overhang at the 3′ end). Supplementary **Table S17** contains the specific oligonucleotides and assembly junctions used for the library generation. The oligos were phosphorylated and annealed to form dsDNA with a 5′ and 3′ overhang. The steps of phosphorylation and annealing were combined and conducted in one pot, by adding 8,000 fmol of the oPool and 1 µl T4 polynucleotide kinase (10 units) to 5 µl 10x T4 ligation buffer and then, adding water until reaching a final volume of 50 µl. After mixing briefly by pipetting the mix was incubated at 37°C for 30 minutes in a thermocycler and then incubated at 65°C for 20 minutes in a thermocycler to heat-inactivate the kinase. For the annealing of the forward and reverse oligo pairs, the following thermocycler steps were added: 95°C for 5 min, 94°C for 15 s, decrease by 1°C, and hold for 30 seconds for 79 cycles. For integrating the dsDNA inserts into DC512, 400 fmol of the dsDNA, 20 fmol of backbone plasmid, 0.5 µL of T4 ligase (1000 units), and 1.5 µL of BbsI (15 units) were added to 2 µL of 10x T4 ligation buffer, then water was added to reach a total volume of 20 µl. A thermocycler was used to perform 35 cycles of digestion and ligation (37 °C for 2 min, 16 °C for 5 min) followed by a final digestion step (60 °C for 10 min) and a heat inactivation step (80 °C for 10 min).

After NdeI digestion (37°C, 1h) of the ligation mix to remove any remaining original backbone plasmids and subsequent ethanol precipitation, 10 µl of the ligation mix was transformed into electrocompetent *E. coli* NEB10 beta (NEB, Ipswich, MA, USA), following the manufacturer’s instructions. After transformation and recovery in 1 ml SOC for 1 h at 37 °C with shaking at 250 rpm, different dilutions of the recovered cells were plated on LB agar containing the appropriate antibiotic and incubated for 16 h to check the number and color of the resulting colonies (ensuring a ∼58X coverage). The rest of the recovered culture was added to 100 mL LB medium containing the appropriate antibiotic and incubated at 37 °C with shaking at 250 rpm to OD_600_ ≈1. Cells were harvested by centrifugation and subjected to plasmid extraction. Sanger sequencing was used to validate the library plasmid DNA.

### Purine screen experiment

*E. coli* strain MG1655 was initially transformed with a dCas9 encoding plasmid (2.0 kV, 200 Omega, and 25 μF). The resulting strain SG332 was then transformed with the sgRNA library by electroporation and recovered in 900 µl SOC for 1.5h at 37 °C with shaking at 250 rpm. Different dilutions of the recovered cells were plated on LB agar containing the appropriate antibiotics and incubated for 16 h to check the number of the resulting colonies (∼56^5^ colonies). The recovered culture was back-diluted to OD_600_ 0.01 in LB medium with appropriate antibiotics and incubated at 37 °C with shaking for 13h. Subsequently, 5 mL of the culture was sampled and the library was extracted by miniprep (Nucleospin Plasmid, Macherey-Nagel) to obtain the initial sgRNA distribution. The calculated amount of culture to reach OD_600_ 0.01 in 50 ml M9 minimal medium, was sampled and washed twice with M9 minimal medium to remove traces of the LB medium. The culture was incubated at 37°C with shaking until it reached OD_600_ 1, allowing ∼6 replications. 5 ml of the culture was sampled at OD_600_ 0.2 and OD_600_ 0.6, and at OD_600_ 1, and the library was extracted by miniprep. The experiment was performed in duplicate starting from two independent transformations of MG1655 with the plasmid library.

### Library sequencing

The sequencing library was generated using the KAPA HiFi HotStart Library Amplification Kit for Illumina® platforms (Roche) and the primers listed in Supplementary Table S17. The first PCR adds the first index. The second PCR adds the second index and flow cell-binding sequence. The amplicons of the first and second PCR reactions were purified using solid-phase reversible immobilization beads (AMPure XP, Beckman Coulter) following the manufacturer’s instructions to remove excess primers and possible primer dimers. The sequencing library samples, with the required DNA concentrations ranging from 100 pg - 200 ng in a total volume of 10 µL, were submitted to the HZI NGS sequencing facility (Braunschweig, Germany) for paired-end 2 × 50 bp deep sequencing with 800,000 reads per sample on a NovaSeq 6000 sequencer.

### Sequencing data processing

Paired-end reads were merged using BBMerge (version 38.69) [52] with parameters “qtrim2=t, ecco, trimq=20, -Xmx1g”. Merged reads with perfect matches were assigned to the gRNA library using a Python script. After filtering guides for at least 1 count per million in at least 4 samples, read counts of each gRNA were normalized by factors derived from non-targeting guides using the trimmed mean of M-values method in edgeR (version 3.28.0) [53]. An extra column was added to the design matrix to capture batch effects between the two replicate experiments. Differential abundance (logFC) of gRNAs between time points and the input library was estimated using edgeR, and a quasi-likelihood F-test was used to test for significance after fitting in a generalized linear model. Spearman correlation between the logFC and predicted scores was calculated for each gene in the purine biosynthesis pathway. For the percentage of efficient gRNAs in predicted efficient gRNAs, gRNAs were first ranked for each gene based on the measured logFC at each time point and the most strongly depleted 20% of gRNAs were considered efficient. Predicted values from each model were also ranked for each gene and the top 20% were considered to be predicted efficient, followed by the calculation of the enrichment of efficient gRNAs in the predicted efficient gRNAs. For the positive predictive value calculation, gRNAs with logFC values within N-fold of the maximum fold change value in each gene were classified as true positives, while the best 3 to 5 predicted gRNAs were defined as predicted positives. The positive predictive values were calculated with the formula PPV = TP / (TP + FP) for all gRNAs at each time point for each fold (N=1.5 - 5 with step of 0.5, TP = True positive, FP = False positive).

### Scoring functions from previous studies

Adapted Python scripts (gRNADesigner.py and DeepSpCas9.py on GitHub) from the source codes of gRNA Designer and DeepSpCas9 were used to calculate the predicted scores. The source code of TUSCAN (https://github.com/BauerLab/TUSCAN) was directly used. For SSC, we used the web-based application at http://crispr.dfci.harvard.edu/SSC/ with the option CRISPR inhibition or activation and 20nt gRNA length. For the Pasteur model, the trained LASSO model was saved from the implemented Python script (Pasteur_model.py on GitHub) based on the jupyter notebook available on GitLab (https://gitlab.pasteur.fr/dbikard/ecowg1).

### Code and data availability

All code necessary to reproduce the results in the manuscript are available at: https://github.com/BarquistLab/CRISPRi_guide_efficiency_bacteria. Raw sequencing data for the CRISPRi purine screen has been deposited in GEO under accession GSE196911. A webserver implementation of the final MERF model is available at: http://ciao.helmholtz-hiri.de.

